# Limited transmission of microbial species among coral reef fishes from the Great Barrier Reef, Australia

**DOI:** 10.1101/2024.02.24.581894

**Authors:** Vincenzo A. Costa, David R. Bellwood, Jonathon C.O. Mifsud, Jemma L. Geoghegan, Erin Harvey, Edward C. Holmes

**Affiliations:** Sydney Institute for Infectious Diseases, School of Medical Sciences, The University of Sydney, Sydney, NSW 2006, Australia; Research Hub for Coral Reef Ecosystem Functions, College of Science and Engineering, James Cook University, Townsville, QLD 4811, Australia; Department of Microbiology and Immunology, University of Otago, Dunedin, New Zealand; Institute of Environmental Science and Research, Kenepuru, Porirua, 5022, New Zealand

## Abstract

Reef fishes account for one-third of all extant marine fishes and exhibit enormous biodiversity within a highly interactive ecosystem. Yet relatively little is known about the diversity and evolution of microbial species (bacteria, viruses, and eukaryotes) associated with reef fish, even though this may provide valuable insights into the factors that shape microbial communities within vertebrate hosts as well as the extent and pattern of cross-species transmission. Through metatranscriptomic sequencing we characterised the viruses, bacteria, and single-celled eukaryotes from 128 reef fish species inhabiting Lizard Island and Orpheus Island on the Great Barrier Reef, Australia. We assessed whether microbial communities differed between islands that are separated by approximately 450 kilometres, and to what extent viruses were able to emerge in new hosts. Notably, despite strong ecological interactions in the reef environment, and the presence of the same families and subfamilies of viruses and bacteria on both islands, there was minimal evidence for the cross-species transmission of individual microorganisms among fish species. An exception was the high prevalence of the bacterial pathogen *Photobacterium damselae* among apparently healthy cardinalfishes from both islands, indicating that these fish species are natural reservoirs within the reef system. Overall, these data suggest that reef fishes have microbial-host associations that arose prior to the formation of the Great Barrier Reef, leading to strong host barriers to cross-species microbial transmission even within a highly interactive and species-rich environment.

## Introduction

Symbiotic interactions are ubiquitous in nature and play an important role in animal, plant, and microbial evolution. In reef systems, the interdependence of corals, fishes, and their microbial symbionts supports fundamental ecological processes, and their dissociation may cause devastating declines in species abundance and reef functioning (1). Healthy tropical coral reefs are characterised by remarkable biodiversity and exceptionally complex interactions. This provides an ideal forum for investigating the evolutionary and ecological factors shaping microbial communities, especially among teleost fishes. Teleost fishes are the most speciose group of vertebrates on coral reefs and rank among the most phylogenetically and ecologically diverse group of vertebrates, accounting for one-third of all currently described marine fishes (2, 3). Reef fishes exhibit exceptional dispersal capabilities. Most have geographic ranges spanning thousands of kilometres, with some species spanning approximately two thirds of the global tropics (4). This level of interconnectivity is also exhibited on individual reefs, with highly complex food webs. Reef fishes display diverse trophic guilds (e.g., carnivores, mobile invertivores, omnivores, planktivores, sessile invertivores, herbivores/detritivores), occur in a range of habitats (e.g., coral, sand, rubble, caves), and commonly live in exceptionally close proximity (5–7).

This high degree of interaction is exemplified in the cryptobenthic reef fishes: behaviourally cryptic species with adult body sizes of approximately 5 cm or less that typically occupy the benthic zone (6). Cryptobenthic reef fishes engage with larger reef fishes through extensive predator-prey interactions, and their frequent consumption is an integral component of coral reef food webs via the transfer of energy from microscopic prey to large predators (8). For example, the dwarf goby (*Eviota sigillata*) has a maximum lifespan of just 59 days and experiences mortality rates of 7.8% per day (9). In addition to predation, fishes interact through cleaning (including the removal of blood-sucking parasites) (10) and extensive food webs of coprophagy (consuming the faeces of other fishes) (11). The potential for microbial transmission on reefs is therefore considerable.

Although reef fish co-exist in a highly diverse and interactive ecosystem, relatively little is known about whether and how microbial composition (i.e., of viruses, bacteria, eukaryotes) differs across fish groups within and among communities. Studies of microbial ecology in teleosts have largely focused on animals utilised in aquaculture and in laboratory model species, with very few investigations of wild ecosystems (12–15). Fish-microbe interactions are highly beneficial for fish nutrition and immunity and are shaped by host and environmental factors including trophic level, age, water quality, and host phylogeny (12, 16–20).

The extent of phylogenetic divergence between animal species has an important impact on virus ecology and evolution, particularly the likelihood of successful cross-species virus transmission, and it is therefore a key determinant of infectious disease emergence (21–25). The “phylogenetic distance” theory posits that microorganisms are more likely to be transmitted between closely related species that have conserved cellular properties, such as cell receptors (26). This idea is supported by numerous studies across a broad spectrum of host taxa (e.g., vertebrates, invertebrates) and pathogen groups including virus-host interactions in reef fishes (16, 22). For example, recent work has shown that reef fishes from a spatially restricted (100 square metre) community from Orpheus Island in the Great Barrier Reef (GBR), Australia, harbour diverse viral assemblages that are highly host-specific despite ample opportunity for cross-species transmission (16).

With approximately 2500 coral reefs and 900 islands, fishes of the GBR constitute a natural model system to investigate spatial patterns of microbial evolution and diversity in vertebrate hosts. Molecular and fossil evidence indicates that the majority of reef fish families originated during the Paleocene and Eocene, approximately 66 to 50 million years ago (Ma). Subsequently, a notable acceleration in lineage diversification took place during the Oligocene and Miocene (34–5.3 Ma), with this shift occurring in the Indo-Australia Archipelago (IAA) (i.e., the IAA biodiversity hotspot) (27). During the Miocene (23-5.3 Ma), the reciprocal diversification of fish and coral species likely led to the development of the functional reef ecosystems that are observable today. By the early stages of the Pleistocene (5.3-0 Ma), almost all reef fish taxonomic groups were established and began their settlement on reefs across all tropical oceans (27), with the formation of the GBR fish communities likely occurring within the last 10,000 years following the stabilization of sea level to its current height ∼6000-8000 years ago (5, 28). Whether and how these colonization events have shaped microbial evolution and diversity is unknown. For example, many reef fishes— particularly cryptobenthics that have limited dispersal—exhibit strong site fidelity, maintaining the same community composition year-round (29). The seemingly consistent community composition might therefore lead to the generation of distinct microbial communities in different geographic areas, that will be most pronounced for rapidly evolving RNA viruses. Conversely, it is possible that the high ecological similarities among disjunct reef locations might result in broadly similar microbial compositions among fish communities.

We characterised the total assemblage of viruses, bacteria, and single-celled eukaryotes from 128 reef fish species spread across Lizard Island and Orpheus Island in the GBR. This study comprised 28 reef fish families, making it one of the largest investigations of microbial diversity and evolution in reef fish undertaken to date. Using metatranscriptomics, we aimed to determine the relationship between host community diversity and microbial diversity and identify whether these communities differ between fish species from two islands separated by approximately 450 kilometres. We also aimed to determine the impact of host ecology on microbial diversity and evolution and to identify whether particular fish groups are potential reservoirs for important viral or bacterial pathogens. This is of particular importance given the high utilisation of reef fish in aquaculture as well as the ongoing threats of biodiversity loss on coral reefs from anthropogenic climate change, itself a significant contributor to the emergence of infectious diseases (30).

## Results

### Composition of sequence reads

We sequenced a total of 10.7 billion RNA reads, including 4.7 billion reads newly generated from Lizard Island fishes. The remaining reads (Orpheus Island) are available on NCBI Sequence Read Archive (SRA) under BioProject PRJNA841039 (16). These data were generated from a total of 140 sequencing libraries, representing 128 reef fish species and 28 families (Figure 1) (mean 54,617,146 reads per library). Fish RNA accounted for 93% of the total reads, followed by RNA associated with cnidarians (5.7%), bacteria (0.31%), single-celled eukaryotes (0.29%), arthropods (0.23%), platyhelminths (0.18%), molluscs (0.04%), poriferans (0.03%), annelids (0.02%), nematodes (0.01%), fungi (0.009%), and viruses (0.009%) (Supplementary Table 1).

**Figure 1.**
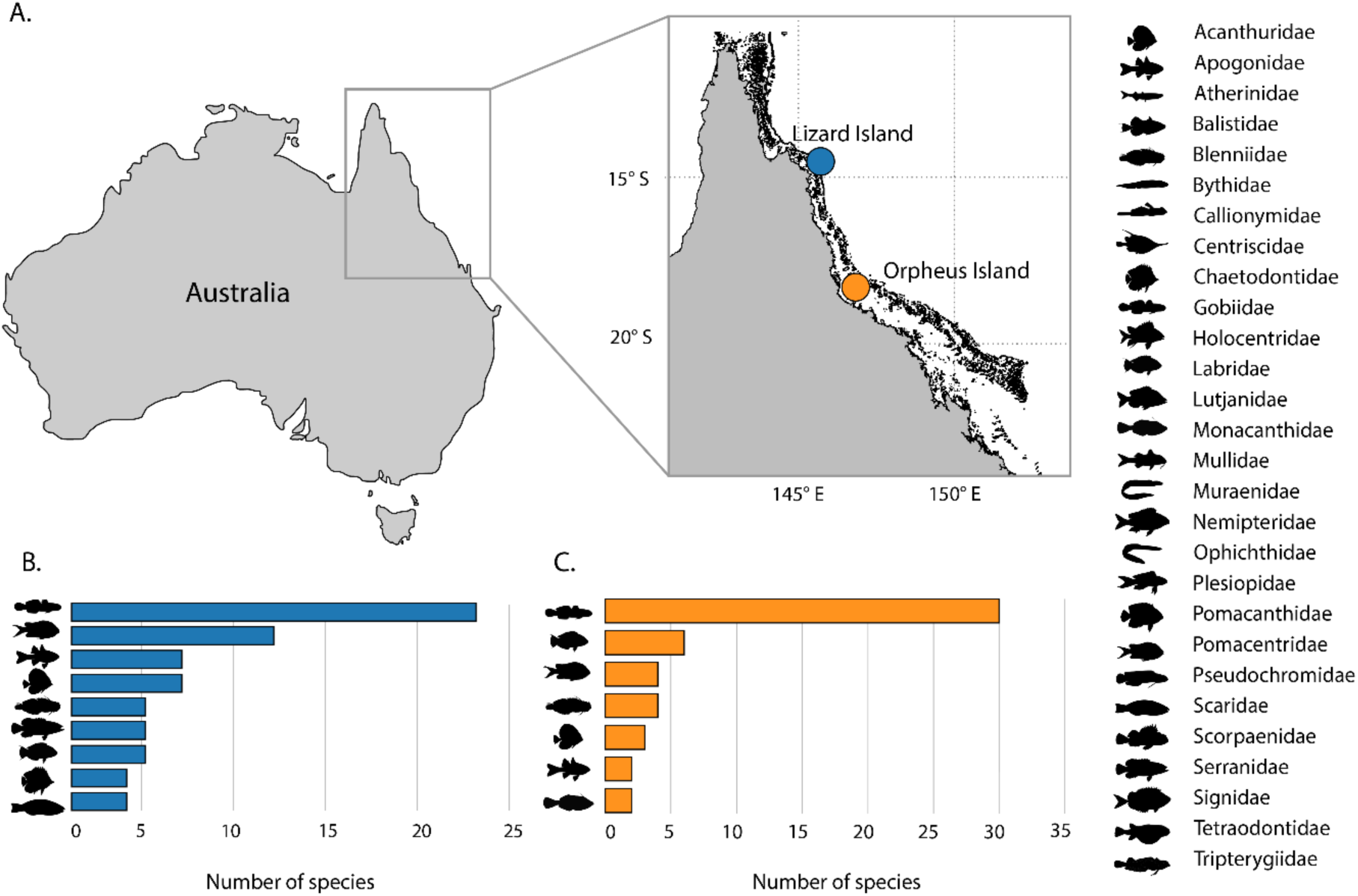
Sampling locations and taxonomic diversity of reef fish. (A) Location of Lizard Island and Orpheus Island in the GBR. (B) Taxonomy of samples collected from Lizard Island. (C) Taxonomy of samples collected from Orpheus Island. All other families with less than one species sampled are omitted from panels B and C.

### Diversity and abundance of the reef fish virome

We identified sequences representing 64 vertebrate-associated viruses (i.e., those likely infecting fish tissues), including 27 newly discovered from Lizard Island fishes (Supplementary Table 2). The *Astroviridae* comprised 32.4% of vertebrate-associated viral reads, followed by the *Iridoviridae* (21.8%), *Picornaviridae* (18.6%), *Chuviridae* (15.9%), *Parvoviridae* (6.8%), *Hantaviridae* (2.9%) with all other groups representing <1% of the total viral reads: *Flaviviridae*, *Orthomyxoviridae (order Articulavirales)*, *Poxviridae*, *Paramyxoviridae*, *Reoviridae*, *Circoviridae*, *Coronaviridae*, *Hepeviridae*, *Rhabdoviridae*, and *Caliciviridae* (Figure 2).

**Figure 2.**
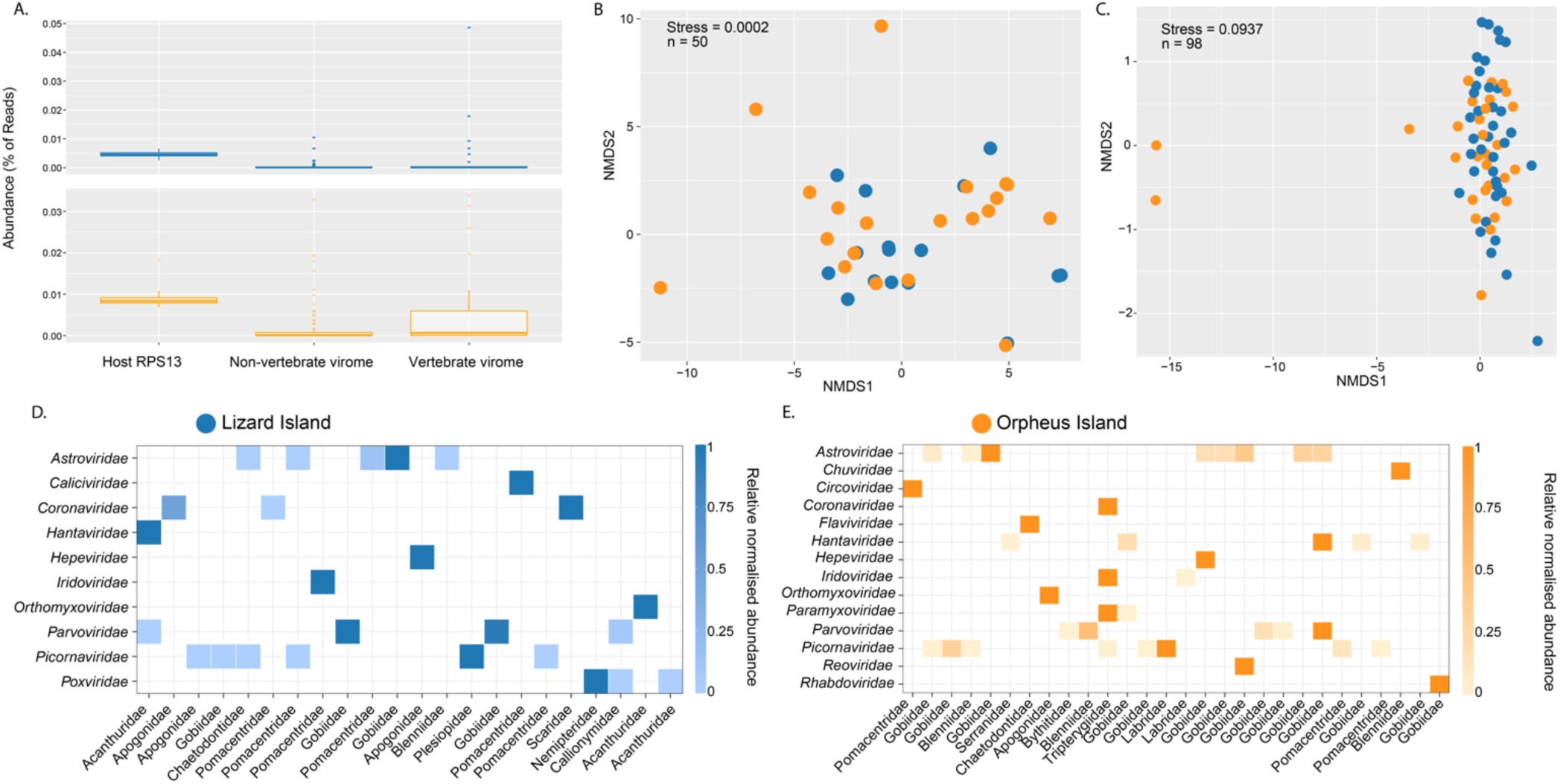
(A) Abundance of host gene marker (RPS13), non-vertebrate virome and vertebrate virome. (B) Non-metric multi-dimensional scaling (NMDS) plot (Bray-Curtis dissimilarity matrix) for vertebrate-associated viruses for each fish species. (C) NMDS plot for non-vertebrate-associated viruses for each fish species. (D-E) Relative normalised abundance of vertebrate-associated viral families from each island.

As well as vertebrate-associated viruses, we identified 194 viruses that were likely infecting porifera, arthropods, molluscs, fungi, plants, and microbial eukaryotes (e.g., dinoflagellates), including 100 that were newly discovered at Lizard Island. Because these viruses were likely associated with fish diet and not infecting the fish themselves, we assume that their presence does not reflect key aspects of fish biology (i.e., immunity or receptor binding) and hence effectively serve as a “negative control” in comparison to the vertebrate-associated viruses. We refer to this group as “non-vertebrate” viruses. The most abundant viral groups in this category were the *Flaviviridae* (24% of non-vertebrate viral reads), unclassified picornaviruses (i.e. “picorna-like” viruses) (22.5%) *Narnaviridae* (21.9%), *Nodaviridae* (8.9%), *Hepeviridae* (6.2%), *Partitiviridae* (4%), *Solemoviridae* (3.2%), *Negevirus* (2.1%) and *Totiviridae* (2%), with all other groups comprising <1%: *Reovirales*, *Weivirus*, *Qinviridae*, *Iflaviridae*, *Dicistroviridae*, *Rhabdoviridae*, *Picobirnaviridae*, *Quenyavirus*, *Bunyavirales*, *Chuviridae*, and *Tombusviridae*.

### Spatial comparisons of the reef fish virome

To determine whether viral composition (i.e., virus families or subfamilies) differed between Orpheus and Lizard islands, we analysed beta diversity using permutational multivariate analysis of variance (PERMANOVA) with the Bray–Curtis dissimilarity matrix. This revealed no significant difference in vertebrate-associated viral communities on both islands (F = 1.42, *p* = 0.06), with overlapping viromes at the viral family/subfamily level (Figure 2b). Accordingly, both islands contained viruses assigned to the *Picornaviridae*, *Astroviridae*, *Parvoviridae*, *Hantaviridae*, *Orthomyxoviridae*, *Coronaviridae*, *Hepeviridae* and *Iridoviridae* (Figure 2d-e). A similar pattern was observed when analysing the non-vertebrate virome (F = 1.53, *p* = 0.07) (Figure 2c) with both islands dominated by unclassified picornaviruses, *Narnaviridae*, *Totiviridae*, *Partitiviridae*, and *Nodaviridae*.

### Evolutionary history and biogeographical patterns of vertebrate-associated viruses

While our analysis revealed similarities at the level of virus family/subfamily, almost all of the viruses identified exhibited levels of genetic divergence that reflected long-term virus-host associations, rather than recent cross-species transmission within each reef ecosystem. The only instance of the same virus being shared between fish species was the presence of highly similar astroviruses (∼96 similarity across the entire genome) in gobies from Orpheus Island (see ref. 16 for full description of these viruses). No viruses were shared among the fishes sampled from Lizard Island.

We now describe the phylogenetic relationships of the shared viral groups in turn. We focus primarily on the relationships of viruses between both islands as well as those newly discovered at Lizard Island.

#### Positive-sense single-stranded RNA viruses (+ssRNA): Picornaviridae, Astroviridae, Hepeviridae, Caliciviridae and, Coronaviridae

The *Picornaviridae* were the most common viral group in our data set with 14 viruses: six from Lizard Island and eight from Orpheus Island. We discovered a group of four novel viruses that formed a distinct clade with the newly formed genus *Danipivirus*, represented by a single virus that is commonly detected in model zebrafishes (31) (Figure 2). In this group it was notable that we identified two relatively closely related viruses (70.6% sequence similarity across the entire polyprotein) in damselfishes (Pomacentridae) from both islands. A more distantly related virus was identified in *Chaetodon baronessa* (Chaetodontidae; 48-49% RdRp similarity with both damselfish picornaviruses) suggesting that these viruses diversified within reef fish, although on an unknown time scale. Evidence of reef diversification was also identified in Pomacentrus nagasakiensis picornavirus and Blenniella picornavirus from Orpheus Island that exhibited 66.1% similarity and were related to fipiviruses found only at Lizard Island (Figure 3).

**Figure 3.**
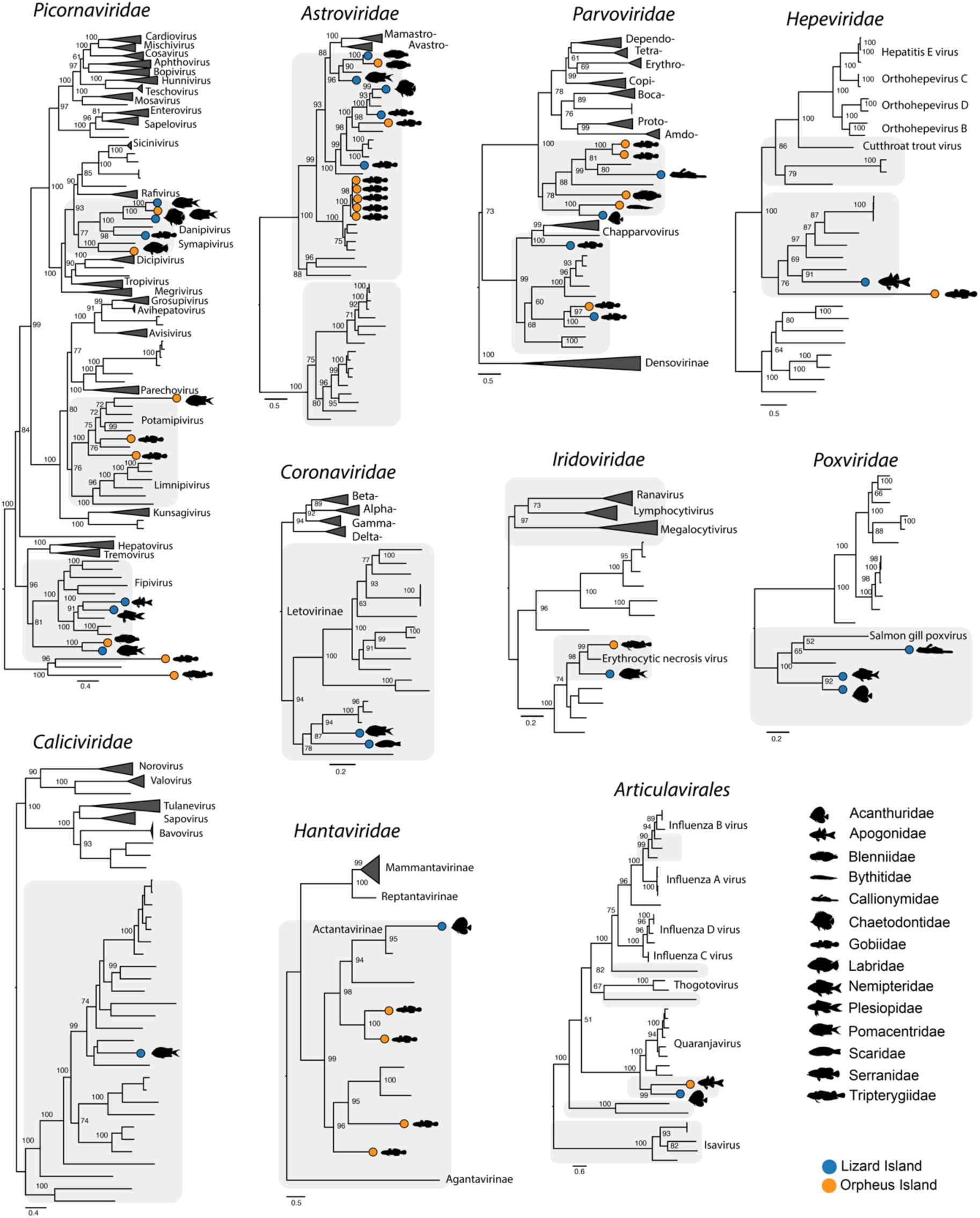
Phylogenetic analysis reveals relationships among reef fish viruses from both islands. Phylogenies were estimated using the RdRp gene for RNA viruses (*Picornaviridae*, *Astroviridae*, *Hepeviridae*, *Hantaviridae, Caliciviridae*, *Coronaviridae*, *Articulavirales*), NS1 gene for parvoviruses and DNA polymerase for iridoviruses and poxviruses. Coloured circles on branch tips represent viruses identified in this study. The scale bar represents the number of amino acid substitutions per site. Shaded branches represent fish viruses. Trees were midpoint rooted for clarity only.

We identified 12 astroviruses, including five novel viruses from Lizard Island. These were spread across three major clades: clade I, represented exclusively by fish viruses including five goby viruses from Orpheus Island (see above); clade II, similarly represented by fish viruses including those from both Lizard and Orpheus Island; and clade III that fell sister to the genera *Mamastrovirus*, found in mammals and *Avastrovirus*, exclusively in birds (Figure 3). Notably, both Plagiotremus tapeinosoma astrovirus (Lizard Island) and Blenniella astrovirus (Orpheus Island) fell within clade III, exhibiting 75.8% sequence similarity in the RdRp gene. Similarly, both Eviota astrovirus and Chaetodon baronessa astrovirus clustered together within clade II, while the other reef fish viruses were more divergent.

Among other +ssRNA viruses, we identified two hepeviruses – Istigobius decoratus hepevirus (Orpheus Island) and Fowleria viaulae hepevirus (Lizard Island) – that grouped with other fish hepeviruses in phylogenetic trees, as well as a novel calicivirus in *Pomacentrus brachialis* that similarly fell within a broad group of fish and amphibian caliciviruses (Figure 3).

Both letoviruses (*Coronaviridae*, subfamily *Letovirinae*) —Chromis atripectoralis letovirus and Scarus psittacus letovirus—from Lizard Island formed a basal clade with Microhyla letovirus and three other viruses identified in African cichlids (32, 33). Together, this group likely formed a novel genus within the *Letovirinae* that includes diverse ectothermic hosts such as amphibians, jawless and ray-finned fishes (32).

#### Negative-sense single-stranded RNA viruses (-ssRNA): Hantaviridae *and* Articulavirales

A notable observation from our previous sampling of reef fish was the detection of hantaviruses in four different goby species at Orpheus Island. In contrast, at Lizard Island we only detected one hantavirus in a surgeonfish (*Acanthurus nigrofuscus*) that was sister to Wenling red spikefish hantavirus (NCBI/GenBank accession: AVM87662.1) and was highly divergent to those found at Orpheus Island (∼29% RdRp similarity). Overall, these viruses fell within the subfamily *Actantavirinae*, that exclusively infects ray-finned fishes. Similarly, we detected two viruses with high levels of divergence (40% similarity) from both islands that were related to quaranjaviruses (order *Articulavirales*) (Figure 3).

#### DNA viruses

We identified reef fish from three lineages within the *Parvoviridae*: the *Parvovirinae*, *Chapparvovirus*, and *Ichthamaparvovirus* groups. Reef fish *Parvovirinae* fell as a sister lineage to mammalian and avian viruses (e.g. *Dependoparvovirus*, *Aveparvovirus*), again with high sequence divergence between islands. For example, the closest relatives among both islands were Acanthurus nigrofuscus parvovirus (Lizard Island) and Dinematichthys parvovirus (Orpheus Island) that shared 62% NS1 gene sequence similarity (Figure 3). Notably, we identified a novel chapparvovirus in *Cryptocentrus strigilliceps* that was related to tilapia parvovirus—a pathogen in farmed Tilapia in China—making it the second fish virus identified in this group (34). Among the genus *Ichthamaparovirus*, both pleurosicya icthamaparvovirus (Lizard Island) and Luposicya lupus ichthamaparvovirus (Orpheus Island) shared a common ancestor (57.9% NSI similarity) and formed a distinct clade with other fish-infecting parvoviruses (16, 33, 35).

Similar patterns were observed in reef fish *Betairidovirinae* (*Iridoviridae*), between both Chrysiptera rollandi iridovirus (Lizard Island) and Enneapterygius tutuilae iridovirus (Orpheus Island). These viruses exhibited 82.2% similarity in the conserved major capsid protein (MCP) and clustered with erythrocytic necrosis virus (ENV).

Poxviruses were only identified at Lizard Island, with phylogenetic analysis placing them with other fish viruses—Salmon gill pox virus and carp edema virus—within the subfamily *Chordopoxvirinae*. Notably, Zebrasoma veliferum poxvirus and Scolopsis bilineata poxvirus shared a common ancestor (80.3% similarity), strongly suggestive of an origin in reef fish (Figure 3).

### Close phylogenetic relationships of non-vertebrate-associated viruses

A common observation in our data set was that genetically diverse reef fish exhibited very similar assemblages of non-vertebrate-associated viruses (Figure 2). This pattern sits in marked contrast to the vertebrate-associated viruses, that were rarely transmitted among species. In particular, we identified the same virus (i.e., 98-99% similarity) in multiple host species within the following groups: unclassified picornaviruses, *Tombusviridae*, *Totiviridae*, *Nodaviridae*, *Narnaviridae*, *Bunyavirales* and (Figure 4). As expected, most of these cases of virus sharing occurred within each island. For example, we identified the same tombusvirus in two blennies and one wrasse from Lizard Island as well as the same narnavirus in three gobies from Orpheus Island (Figure 4). A notable exception was the presence of two nodaviruses identified in *Fowleria vaiulae* from both islands that exhibited 98% similarity in the RdRp gene. While nodaviruses are capable of infecting fish species (e.g. nervous necrosis virus) we detected reads associated with decapods in both libraries. These reads, in combination with the phylogenetic positions of these viruses (i.e., highly divergent from nervous necrosis virus and more related to crustacean viruses) strongly implies that these viruses are of dietary origin.

**Figure 4.**
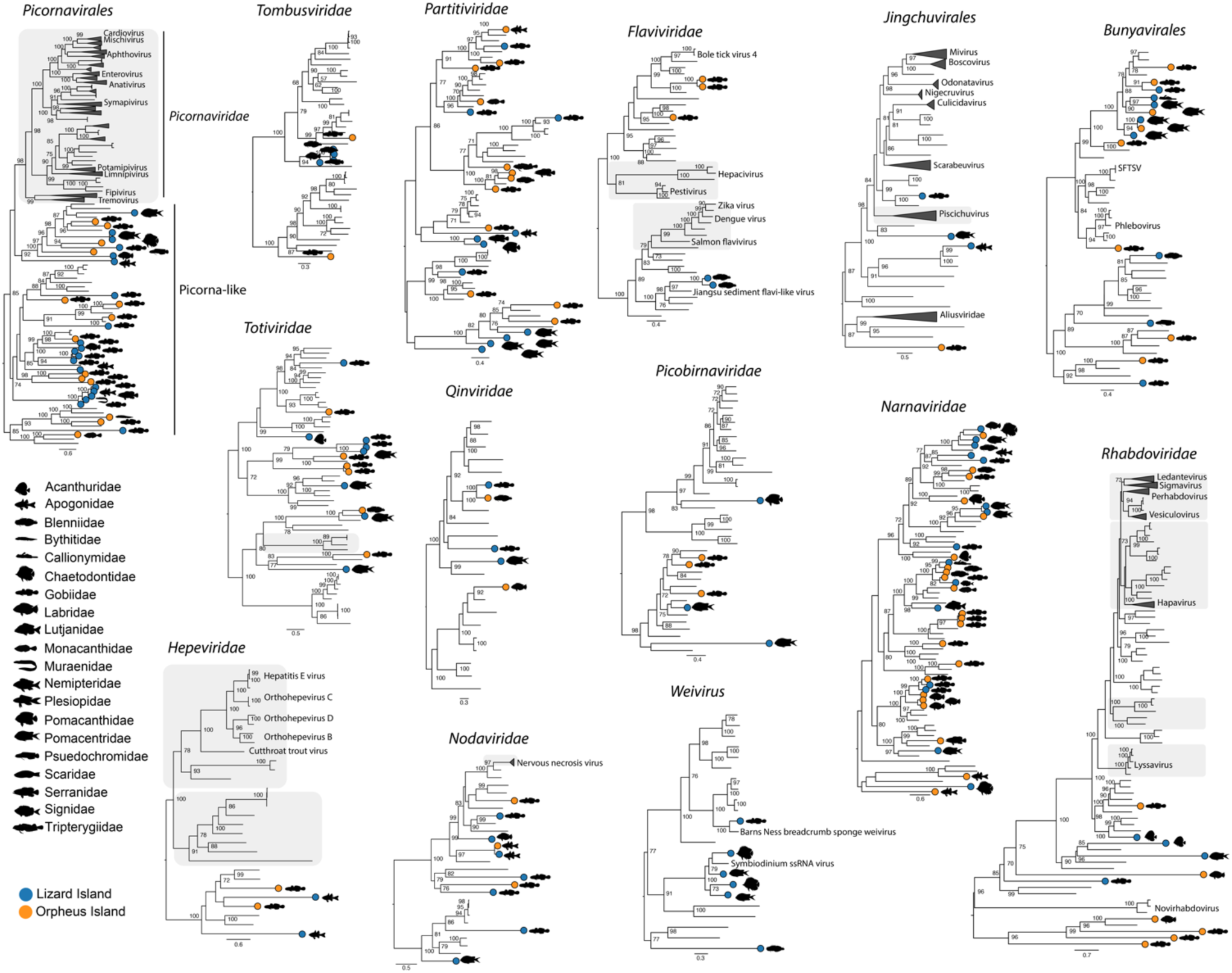
Close phylogenetic relationships among non-vertebrate-associated viruses sampled from reef fish. Phylogenies were estimated using amino acid sequences of the RdRp gene. Coloured circles on branch tips represent viruses identified in this study. The scale bar represents the number of amino acid substitutions per site. Shaded branches represent vertebrate-associated viruses. Trees were midpoint rooted for clarity only.

### Diversity and abundance of bacteria in reef fish assemblages

After the removal of cyanobacteria, the phylum *Proteobacteria* accounted for 73.5% of the total sequence reads from bacteria, followed by *Firmicutes* (9%), *Actinobacteria* (8%), *Bacteroidetes* (2.9%), *Spirochaetes* (2.3%), *Fusobacteria* (1.7%), with all other phyla each representing <1% (Figure 5a). At the family level, the *Vibrionaceae* (42.7%) and *Enterobacteriaceae* (11.6%) were present at the highest frequencies followed by the *Endozoicomonadaceae* (5.9%), *Comamonadaceae* (5.9%), *Shewanellaceae* (5.8%), *Propionibacteriaceae* (5%), *Clostridiaceae* (4.9%), *Micrococcaceae* (2.3%), *Fusobacteriaceae* (2%), *Pseudomonadaceae* (1.3%), and *Mycoplasmataceae* (1.3%) (Figure 5).

**Figure 5.**
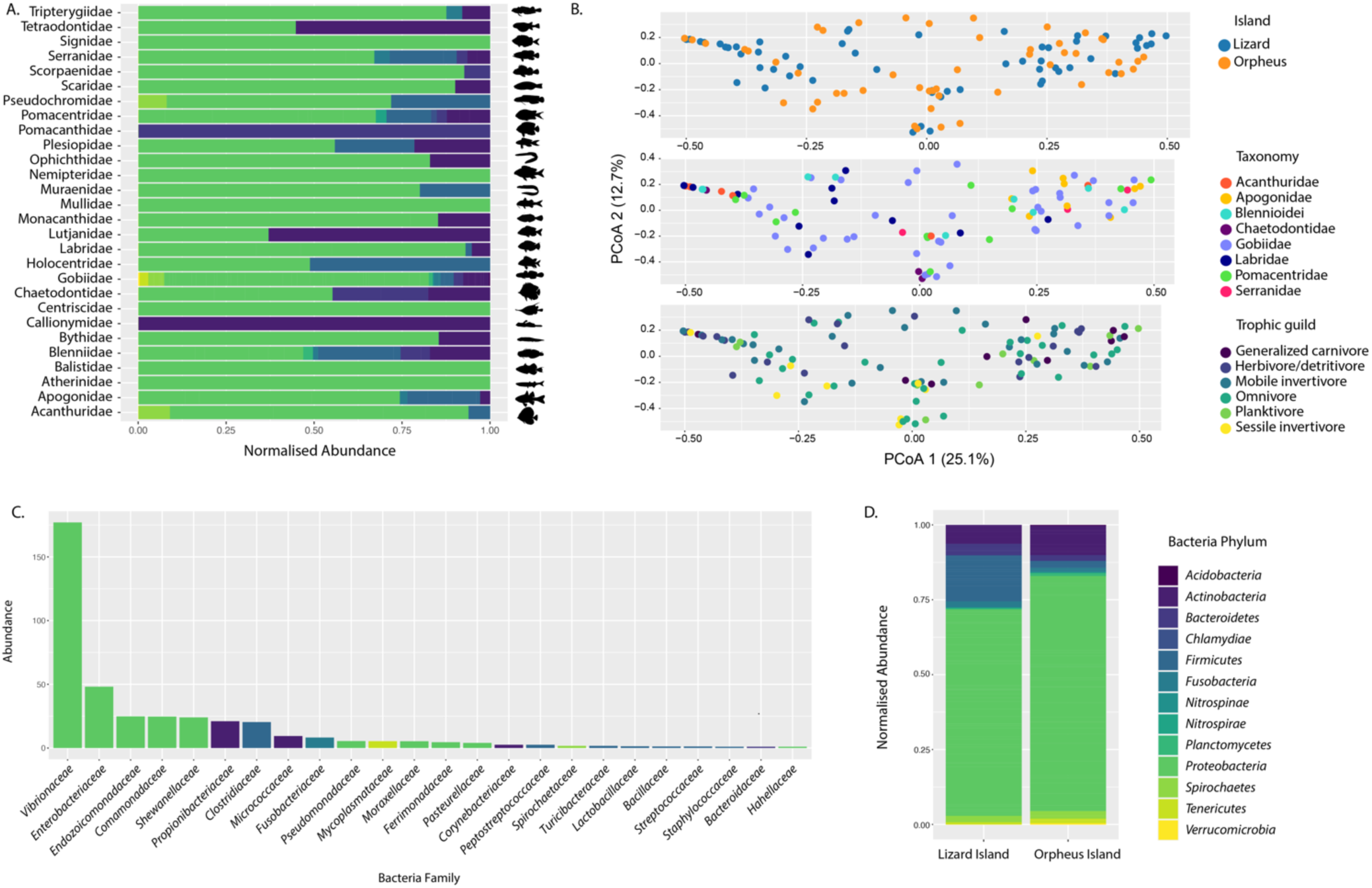
(A) Normalised abundance of bacterial phyla for each reef fish family. (B) Principal coordinate analysis (PCoA) plots of bacterial communities for island, host taxonomy and trophic guild using Bray-Curtis dissimilarity matrix. (C) Abundance of bacterial families coloured by phylum. (D) Normalised abundance of bacterial phyla for each island.

To explore patterns of bacterial diversity between both islands and among reef fish groups, we performed a principal coordinate analysis (PCoA). This revealed no partitioning according to reef location with largely overlapping bacterial communities between the two islands (*F* = 1.711, *R*^2^ = 0.015, *p* = 0.082) (Figure 6). PERMANOVA revealed significant differences in bacterial composition between fish taxonomic groups (*F* = 2.42, *R*^2^ = 0.153, *p* = 0.001). However, there was overlap among all fish species, particularly the gobies, which may be explained by their high involvement in coral reef food webs (5, 6) (Figure 5b).

**Figure 6.**
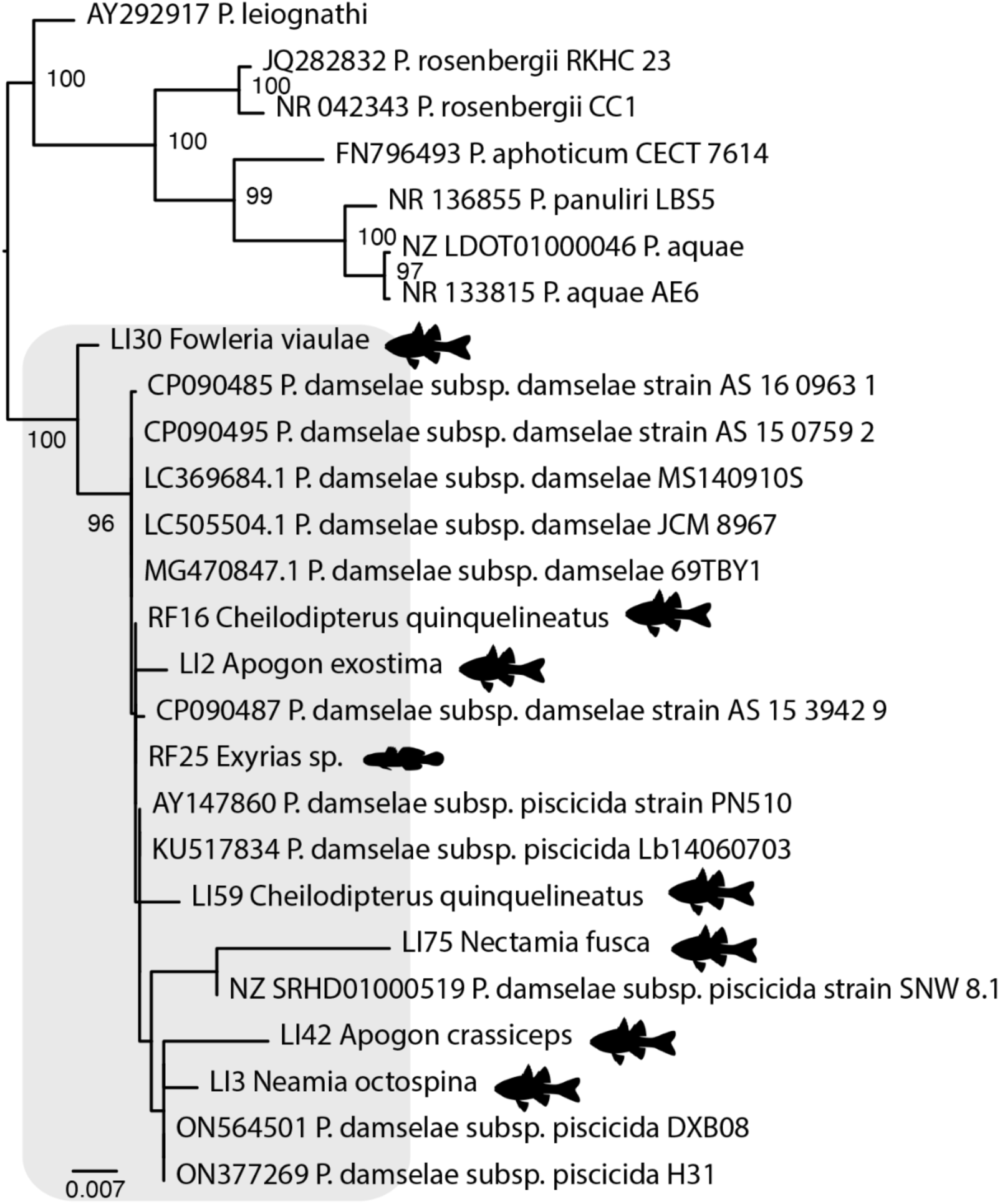
Maximum likelihood phylogeny of the genus *Photobacterium*, estimated using nucleotide sequences of the 16S gene. Fish silhouettes represent individuals identified in this study. The scale bar represents the number of nucleotide substitutions per site. Tree was midpoint rooted for clarity only.

Despite the high degree of overlap in bacterial families among the reef fishes, there was limited evidence for the sharing of individual bacterial species between fish species. Notably, however, we identified *Photobacterium damselae*—an important pathogen in aquaculture— in 88% of the cardinalfish species examined from both islands (Figure 6). We also detected other potentially pathogenic *Vibrionaceae*, including four that were related to those within the *Vibrio harveyi* clade: *V. parahaemolyticus*, *V. campbellii*, and *V. owensii*. (Supplementary Figure 1). In addition, we identified *V. fortis*—an opportunistic pathogen of coral—in the surgeonfish, *Ctenochaetus binotatus* (36) (Supplementary Figure 1).

To assess the impact of fish ecology on bacterial composition, we grouped fish species into six trophic guilds: carnivores, mobile invertivores, omnivores, planktivores, sessile invertivores, herbivores/detritivores (7). While we detected significant differences in bacterial composition between these groups (*F* = 1.485, *R*^2^ = 0.074, *p* = 0.035), there was similarly high overlap with no clear partitioning according to host trophic guild (Figure 5b).

### Composition of single-celled eukaryotes

Finally, we identified transcripts representing single-celled eukaryotes from 15 phyla. Dinoflagellates were the most abundant group (46% of the total eukaryotic reads) followed by Bacillariophyta (21.6%), Foraminifera (12.6%), Cercozoa (5.8%), Euglenozoa (2.9%), Apicomplexa (2.7%), Ciliophora (2.5%), Endomyxa (1.1%), Parabasalia, Haptista, Fornicata, and Heterolobosea (all less than one percent). Fungi—Ascomycota, Basidiomycota, Microsporidia—were identified at much lower frequencies, representing only 3.3% of the total reads in this category.

Using these data, we performed a PCoA based on fish location, taxonomy and ecology. As with the analysis of virome composition, this revealed overlapping microbial communities between both islands (*F* = 1.208, *R*^2^ = 0.015, *p* = 0.181). However, there were significant differences in microbial communities between fish taxonomic groups (*F* = 1.384, *R*^2^ = 0.119, *p* = 0.002), which may be driven by the separation of pomacentrids from other groups, in turn reflecting the larger diversity of microorganisms identified in this group (Figure 7a-b). When assessing the impact of host ecology, we similarly identified substantial overlap with no clear partitioning (Figure 7). Apart from *Symbiodinium* spp., which form symbiotic relationships with coral rather than fish and hence are dietary-associated, we found no evidence for the same microbial species present in multiple fishes. Among the Apicomplexa, which are common parasites of vertebrates, we detected reads associated with *Goussia* spp. in *Chaetodon aureofasciatus*, *Halichores melanurus*, and *Eviota melasma*. Overall, the parasitic families, such as the Eimeriidae (Apicomplexa), Trypanosomatidae, and Ichthyobodonidae (both Euglenzoa) were found in eight libraries at low transcript abundances, representing an average of 0.19% of the reads in each library.

**Figure 7.**
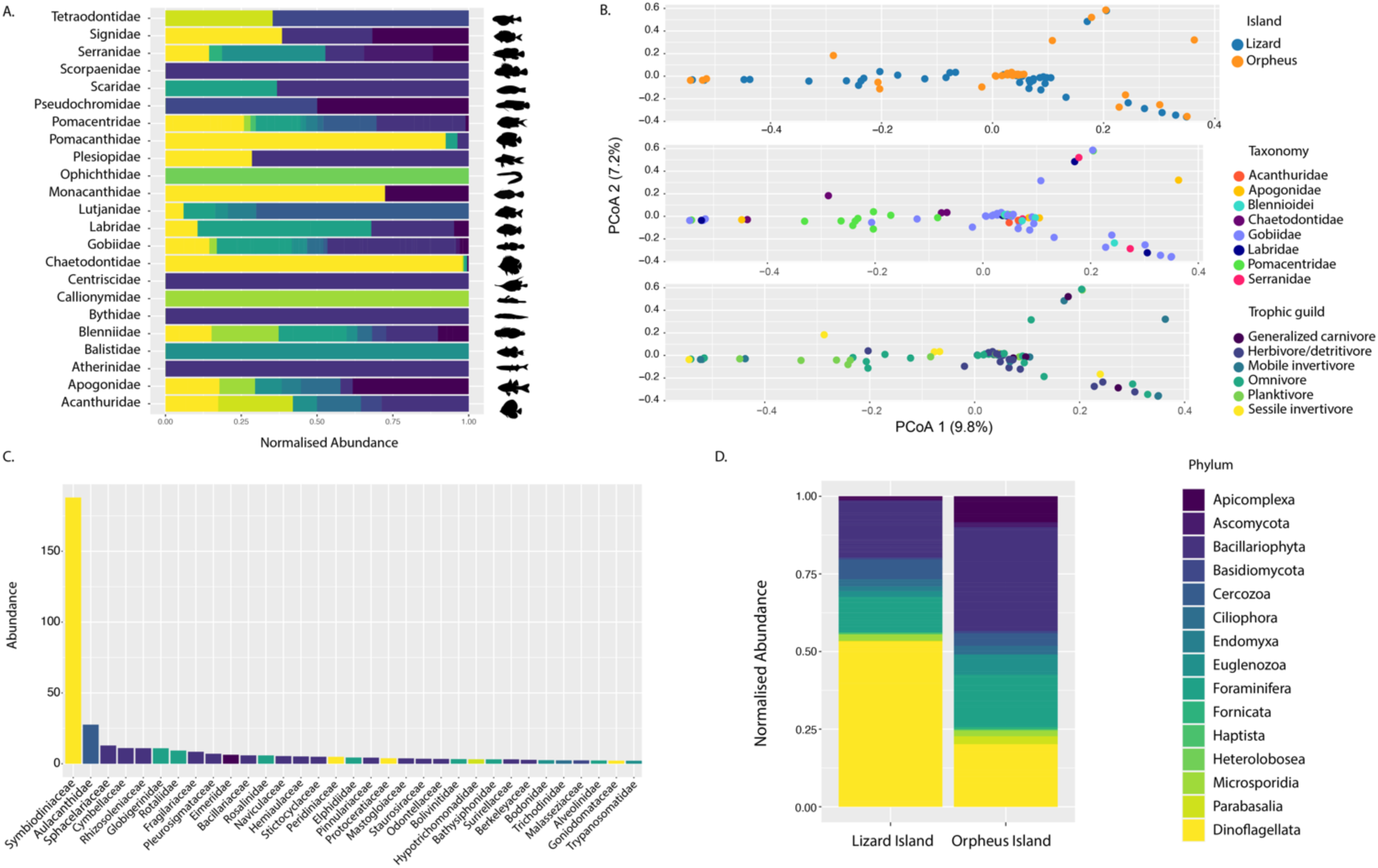
(A) Normalised abundance of single-celled eukaryotic phyla for each reef fish family. (B) Principal coordinate analysis (PCoA) plots of single-celled eukaryotic communities for island, host taxonomy and trophic guild using Bray-Curtis dissimilarity matrix. (C) Abundance of single-celled eukaryotic families coloured by phylum. (D) Normalised abundance of single-celled eukaryotic phyla for each island.

## Discussion

Our metatranscriptomic analysis of 128 reef fish species revealed no significant differences in the composition of viral and microbial families/subfamilies between two islands located ∼450 kilometres apart in the Australian Great Barrier Reef. Fish sampled from islands were associated with the presence of the *Picornaviridae*, *Astroviridae*, *Parvoviridae*, *Hantaviridae*, *Orthomyxoviridae*, *Coronaviridae*, *Hepeviridae* and *Iridoviridae*, as well as *Proteobacteria* of the *Vibrionaceae, Enterobacteriaceae*, and *Endozoicomonadaceae*. Also of note was that within each group of viruses we observed high levels of genetic diversity, with minimal evidence for the same virus being shared among fish species (both within and between islands), despite strong ecological interactions in the reef ecosystem and our spatially restricted sampling.

These findings offer further support for recent studies of virus ecology in fish that show that host phylogeny has a strong influence on virome composition (16, 33). For example, closely related African cichlid species, that have evolved *in situ* within Lake Tanganyika over the last 10 million years, exhibit highly similar viromes with high levels of cross-species transmission and viral generalism (33). African cichlids are members of a single family, the Cichlidae, and exhibit some of the lowest pairwise genetic distances observed between vertebrates (e.g., differences of 0.03% between some species). In contrast, reef fish communities are composed of several divergent families, including 28 examined in this study (7, 37), many of which were established around 66 Ma, occupying the biogeographic region – Tethys – now covered by Europe and the Mediterranean Sea (27). During the Oligocene there was a shift in species richness from Tethys to the IAA, eventually forming the IAA biodiversity hotspot (27, 38). Most reef fish genera formed within the IAA around 30-15 Ma, with accelerated speciation during the Miocene (27).

A notable difference between cichlids and reef fishes, which may explain their strikingly contrasting levels of cross-species transmission, is that cichlids rapidly evolved within Lake Tanganyika, while reef fishes diversified within the IAA, over many millions of years, prior to their settlement on the GBR (27, 39). As the GBR that exists today formed after Pleistocene sea level rises, reef fish communities only became established during the last ∼10,000 years, such that the genetic boundaries inhibiting cross-species transmission were already established in the IAA before their settlement at reef locations. In marked contrast, the adaptive radiation of the African cichlids would have provided a more favourable environment for cross-species transmission as their diversification occurred rapidly within the confinements of Lake Tanganyika, generating a large pool of closely related host species for infection. Indeed, a time-calibrated phylogeny of cichlid hepaciviruses showed that elevated rates of virus diversification coincided with a period of rapid cichlid speciation (33).

These ecological and evolutionary patterns were similarly observed across the bacteriome, with *P. damselae* the only bacterium that was shared among multiple fish species. Indeed, *P. damselae* was primarily identified in cardinalfishes (Figure 6), further illustrating a phylogenetic effect. However, it is likely that this association represents a long-term symbiotic relationship between cardinalfishes and *P. damselae*, particularly as it was detected in *Cheilodipterus quinquelineatus* from both islands, as well as its presence in 88% of the cardinalfish libraries examined. This strongly suggests that *P. damselae* forms part of the natural microbiome in these species. It is noteworthy that most cardinalfish species are nocturnal on coral reefs, and the genus *Photobacterium* is renowned for its bioluminescence—via the expression of *lux* genes—including some strains of *P. damselae* (40). Indeed, *Photobacterium mandapamensis* is a bioluminescent symbiont of the urchin cardinalfish, *Siphamia tubifer*, where it provides light to attract prey (40, 41). Several cardinalfish species have evolved specialized “light organs” that harbour bioluminescent *Photobacterium*; however, these specialized organs are not present in all species (42). It is important to note that we did not observe the expression of *lux* genes in any of our libraries, and these species are not recognized for possessing a bioluminescent system. The presence of *P. damselae* in almost all Apogonidae libraries implies a longstanding relationship between *Photobacterium* and cardinalfishes on coral reefs, suggesting that this genus might have originated and diversified in the reef environment. Overall, these data show that cardinalfishes serve as natural hosts for *P. damselae* and should be monitored closely, particularly if interacting with farmed populations that are often severely affected by *P. damselae* infection (43).

The identification of two iridoviruses that were related to Erythrocytic necrosis virus (ENV; 83-89% MCP similarity) was also noteworthy. ENV is an important pathogen in the North Atlantic and North Pacific oceans (44), and the presence of an ENV-like virus at Lizard Island is consistent with a previous study that identified viral erythrocytic necrosis (VEN) in a juvenile triggerfish (*Rhinecanthus aculeatus*) at this location (45). Moreover, this study identified VEN-like bodies in *R. aculeatus* erythrocytes that were also found in the digestive tract of associated gnathiid isopods that are common blood feeding parasites. Similarly, we detected reads from gnathiids in *C. rollandi* from Lizard Island, implying that gnathids could act as vectors in the marine environment (45), although we did not detect any gnathiid reads in *Enneapterygius tutuilae* from Orpheus Island. This association warrants further investigation and has the potential to improve control measures against pathogenic ENV in susceptible species such as pink (*Oncorhynchus gorbuscha*) and chum (*Oncorhynchus keta*) salmon (44).

It was notable that we identified a low number of protozoan parasites (n = 8), such as apicomplexans and trypanosomes. In a similar manner, we detected low levels of fungal reads (e.g., Ascomycota, Basidiomycota and Microsporidia), supporting the idea that the marine environment, particularly coral reefs, contains low fungal biomass perhaps because of the oligotrophic conditions compared to nutrient-rich terrestrial environments (46). The vast majority of reads in this category came from dinoflagellates, diatoms, and foraminifera that likely came from the reef environment, particularly dinoflagellates that form symbioses with corals (1). Indeed, we discovered six viruses that fell within the “Weivirus” group—a group of dinoflagellate, poriferan and mollusc viruses—grouping with a virus recently identified in *Symbiodinium spp.* (47, 48) (Figure 4).

While our microbial profiling was able to detect diverse microbial communities, it is important to note that our sampling was initially performed for virological analysis (16), such that our methodology was optimized for the detection of viruses (e.g. ribosomal depletion). This may have limited our ability to detect bacteria and microbial eukaryotes. Moreover, there were necessary limitations in our sampling that impacted the power of our statistical analyses. For example, there was a considerably higher number of gobies sampled compared to other reef fish families (Figure 1). Moreover, these comparisons are based on combined tissues, such as liver and gills or whole fish (i.e. cryptobenthic reef fishes). Overall, given that metatranscriptomics is solely based on RNA-sequencing, we were only able to detect microbes and DNA viruses that were expressing genes during the time of sampling.

In summary, while our metatranscriptomic of reef fish communities from islands separated by ∼450 kilometres revealed the same viral and bacterial families across both islands, there was strikingly little evidence for cross-species transmission within reef fish communities. As such, these data support the concept that fish are rich in microbial diversity, but that there are strong barriers to infection in host communities that display high levels of genetic diversity.

## Materials and methods

### Ethics

Fish were collected under a Great Barrier Reef Marine Park Authority permit (G16/37684.1) and James Cook University Animal Ethics permit A2752.

### Fish sample collection

Building on transcriptome data from our previous sampling of reef fish from Orpheus Island (n = 192 fishes; 16 reef fish families) (16), an additional 163 individuals (24 families) were collected at Lizard Island (14°40’08″S 145°27’34″E) during January 2022 (Supplementary Table 3). All fishes were intact with no visible signs of disease, and the vast majority were adults. These animals were captured using an enclosed clove oil method (16) from both Mermaid Cove and the Lagoon entrance, located on the north and south side of Lizard Island (Supplementary Table 1). All fish caught were placed either dissected (liver and gills) or whole in RNAlater and then transported to the lab on ice (Supplementary Table 1). Specimens were stored at −80°C until RNA extraction. Overall, we sampled 1-12 individuals per species (with a mean of 3 per species).

### RNA extraction, metagenomic library preparation and next-generation sequencing

As described previously (16), tissue specimens (e.g. liver and gills or whole fish) were collectively processed as a single extraction for each individual fish sample. The combined tissues were submerged in 600*μ*l of lysis buffer containing 30 *μ*l of foaming reagent (Reagent DX, Qiagen) and 60*μ*l of *β*-mercaptoethanol (Sigma-Aldrich). Tissue samples were homogenized with a TissueRuptor (Qiagen) for up to 1 minute at 5,000 rpm. The homogenate was centrifuged at maximum speed for three minutes to remove tissue residues. The RNA was then extracted from the resulting clear supernatant using the RNeasy Plus Mini Kit (Qiagen, Hilden, Germany), following the manufacturer’s guidelines.

RNA quantification was conducted utilizing a UV–Vis cuvette spectrophotometer (DeNovix, Delaware, USA) and a parallel capillary electrophoresis instrument (Fragment Analyzer; Agilent, CA, USA). RNA from individual fishes were pooled according to species, resulting in 79 RNA sequencing libraries newly generated from Lizard Island. All libraries were prepared using the TruSeq Total RNA Library Preparation Protocol (Illumina). Ribo-Zero Plus Kit (Illumina) was employed for host ribosomal RNA depletion, and paired-end sequencing (150 bp) was performed on the NovaSeq 6000 platform (Illumina). To mitigate index hopping and minimize false virus–host assignments, each library was sequenced on two different lanes. Library construction and metatranscriptomic sequencing were performed by the Australian Genome Research Facility.

### Assembly of reef fish viromes

We replicated the same methodology we used previously (16). Accordingly, raw RNA sequencing reads were quality trimmed using Trimmomatic v.0.38, employing the parameters SLIDINGWINDOW:4:5, LEADING:5, TRAILING:5, and MINLEN:25, and assembled into contigs using MEGAHIT v.1.2.9, with default parameter settings (49, 50). Assembled contigs were compared against the NCBI non-redundant protein (nr) and nucleotide (nt) databases (August 2022) using DIAMOND (BLASTX) (v.2.0.9) and BLASTn (51). To enable the identification of divergent viral sequences, we used an e-value search threshold of 1 x 10^-5^. Contigs with top matches to the kingdom “Viruses” (NCBI taxid: 10239) were predicted as open reading frames (ORFs) using Geneious Prime (v.2022.0) (www.geneious.com) (52). To remove false positives, all putative viral ORFs were translated into amino acid sequences and used as a query to perform a second search (BLASTP) against the NCBI nr database using Geneious Prime. ORFs with top matches to fish genes were deemed as false positives and removed from further analysis. To determine whether our putative viral contigs were expressed endogenous viral elements (EVEs) we screened for disrupted ORFs and flanking host regions using CheckV and BLASTn (16, 33). Viral contig contamination and completion was determined using CheckV (53). Transcript abundances of both host (RPS13 gene) and virus (16, 17, 54) were calculated using RNA-Seq by Expectation Maximization (RSEM) (v.1.3.0) and coverage was assessed by mapping using Bowtie2 (v.2.3.3.1) (55, 56).

### Taxonomic assignment and genome annotation of reef fish viruses

We aligned the amino acid sequences of our putative viruses (partial or complete) with the complete sequences of related viruses available on NCBI/GenBank using the E-INS-i algorithm in MAFFT v.7.450 (57). To determine whether our viruses were novel species, we used levels of sequence similarity and phylogenetic relationships (see below) as specified by the International Committee of Viral Taxonomy (ICTV) (https://talk.ictvonline.org) for each viral genus/family. We used these criteria to determine whether a virus was likely infecting reef fishes (i.e. vertebrate-associated) or of “non-vertebrate” origin, such as those derived from fish diet, microbiome or environment (16, 17, 54). Viral genomes were annotated with the Live Annotate and Predict tool in Geneious using reference sequences from NCBI/GenBank, with a similarity threshold of 20%. We also used the NCBI conserved domain (CDD) search tool and InterProScan with the TIGRFAMs (v.15.0), SFLD (v.4.0), PANTHER (v.15.0), SuperFamily (v.1.75), PROSITE (v.2022_01), CDD (v.3.18), Pfam (v.34.0), SMART (v.7.1), PRINTS (v.42.0), and CATH-Gene3D databases (v.4.3.0) (58).

### Viral phylogenetic analysis

To infer the evolutionary relationships of both the vertebrate and non-vertebrate associated viruses, we aligned the translated contigs with background protein sequences from each viral family/subfamily/genus selected from the ICTV classification and obtained from NCBI/GenBank. For RNA viruses, we used the conserved RNA-dependent RNA polymerase (RdRp), while for DNA viruses we used the DNA polymerase. Amino acid sequence alignments were trimmed using TrimAl (v.1.2) with a gap threshold of 0.9 and a variable conserve value (59). The best-fit model of amino acid substitution was estimated with the “ModelFinder Plus” (-m MFP) flag in IQ-TREE (v.1.6.12) (60, 61). We used a maximum likelihood approach to estimate phylogenetic trees using IQ-TREE, with 1000 bootstrap replicates. Trees were annotated using FigTree (v.1.4.4) (http://tree.bio.ed.ac.uk/software%20/figtree/).

### Virus nomenclature

Viruses were provisionally named (i.e., awaiting ICTV confirmation) according to host species (e.g. *Halichoeres melanurus ranavirus*) as described previously (16).

### Microbial profiling

To screen for transcripts associated with bacteria or single-celled eukaryotes, we aligned our contigs to a custom database comprising all nucleotide sequences available on NCBI (with the removal of environment or artificial sequences) using the KMA aligner and CCMetagen (62, 63). We also used this output to assess metazoan reads—e.g., arthropod, mollusc, platyhelminth, nematode—that may represent potential vectors for virus transmission. For instances in which CCMetagen identified a microbe at the species level, we validated these taxonomic assignments by: (i) performing an additional search (BLASTn) against a custom 16S (bacteria) or 18S (eukaryote) rRNA database, and (ii) analysing the BLASTX output (see above) for hits to bacterial or eukaryotic proteins. The contigs from these BLAST hits were predicted into ORFs, translated into amino acid sequences, and used as a query to perform a second search against the NCBI using BLASTP for further validation. The 16S or 18S rRNA gene was then utilised for phylogenetic analysis as a final validation.

### Analysis of Beta diversity

To compare viral and microbial communities between reef fish assemblages, we calculated beta diversity using a Bray–Curtis distance matrix with the phyloseq package in R (64). The variables assessed were host taxonomy, location (i.e. island) and trophic guild. Accordingly, fish species were categorised into six trophic guilds: carnivores, mobile invertivores, omnivores, planktivores, sessile invertivores, herbivores/detritivores as described in (7). We based our analysis on groups with three or more species. These data were then tested using permutational multivariate analysis of variance (PERMANOVA) with the vegan package (adonis) (65). All plots were constructed using ggplot2 in R (66).

## Data Availability

Raw sequence reads have been deposited in the Sequence Read Archive (NCBI/SRA) under BioProject PRJNA1078998. All viral sequences discovered have been deposited in NCBI/GenBank under the accessions XXXX-XXXX. All phylogenetic trees, tables, and chrona plots are available on GitHub under the repository https://github.com/vcosta16/reeffishvirome.

## Acknowledgements

This work was funded by an Australian Research Council Discovery Project (DP200102351) and a National Health and Medical Research Council Investigator grant (GNT2017197) to ECH, and an ARC Laureate Fellowship (FL 190100062) to DRB. JLG is funded by a New Zealand Royal Society Rutherford Discovery Fellowship (RDF-20-UOO-007) and a Marsden Fund Fast Start (20-UOO-105). We acknowledge the University of Sydney for providing the high-performance computer cluster “Artemis”, that was used for this study.

## Author Contributions

VAC and ECH conceptualised the study. VAC, DRB and JCOM performed the sampling. VAC performed the analyses. VAC wrote and prepared the original draft. VAC, JCOM, ECH, EH, DRB, and JLG edited and revised the manuscript. ECH and JLG funded the project. EH and ECH supervised the project.

