## Supplementary figures and images for "Limited transmission of microbial species among coral reef fishes from the Great Barrier Reef, Australia"

### Supplementary Figure 1

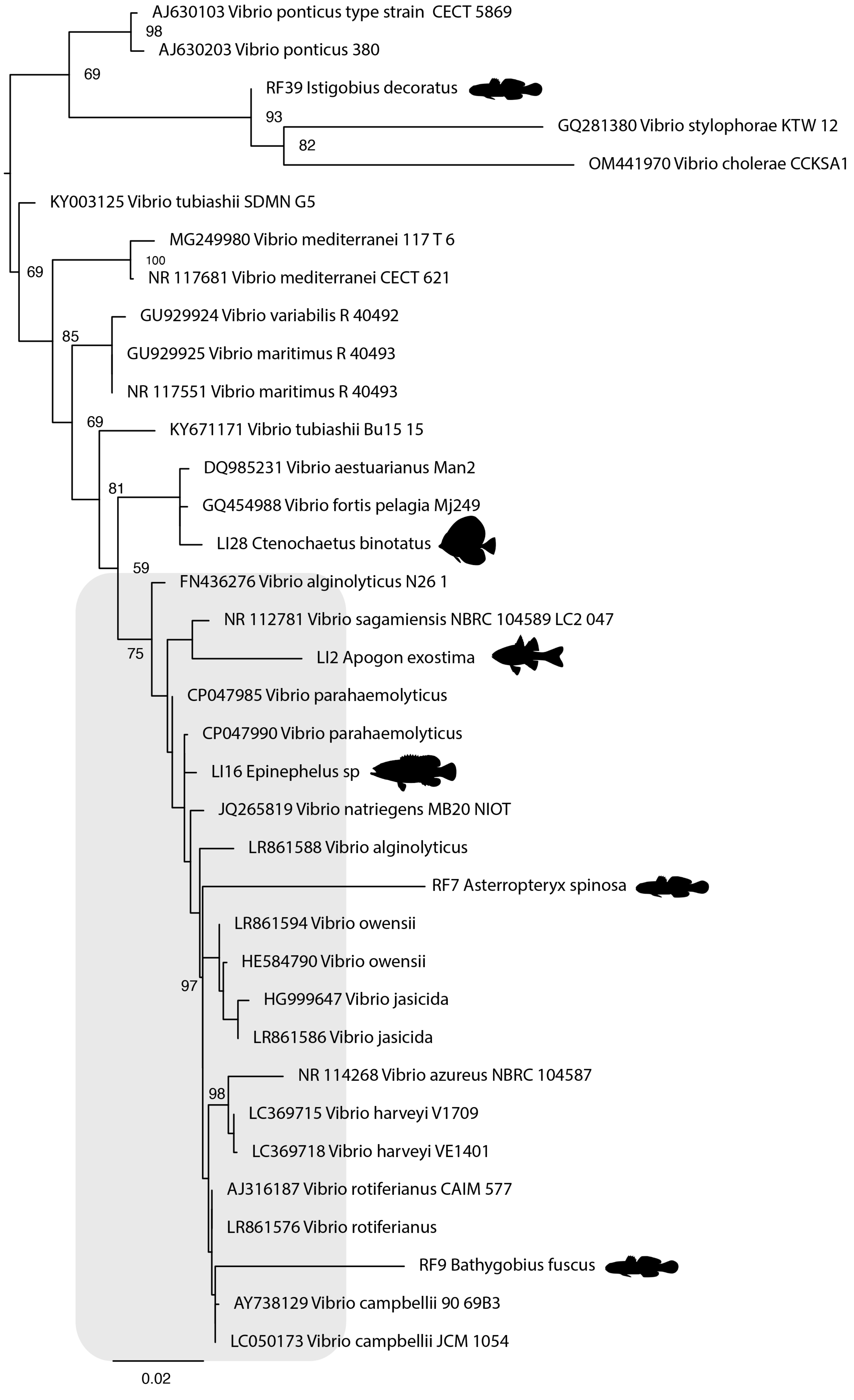
